# Neural tracking of the speech envelope in cochlear implant users

**DOI:** 10.1101/359299

**Authors:** Ben Somers, Eline Verschueren, Tom Francart

## Abstract

**Objective:** When listening to speech, the brain tracks the speech envelope. It is possible to reconstruct this envelope from EEG recordings. However, in people who hear using a cochlear implant (CI), the artifacts caused by electrical stimulation of the auditory nerve contaminate the EEG. This causes the decoder to produce an artifact-dominated reconstruction, which does not reflect the neural signal processing. The objective of this study is to develop and validate a method for assessing the neural tracking of speech envelope in CI users.

**Approach:** To obtain EEG recordings free of stimulus artifacts, the electrical stimulation is periodically interrupted. During these stimulation gaps, artifact-free EEG can be sampled and used to train a linear envelope decoder. Different recording conditions were used to characterize the artifacts and their influence on the envelope reconstruction.

**Main results:** The present study demonstrates for the first time that neural tracking of the speech envelope can be measured in response to ongoing electrical stimulation. The responses were validated to be truly neural and not affected by stimulus artifact.

**Significance:** Besides applications in audiology and neuroscience, the characterization and elimination of stimulus artifacts will enable future EEG studies involving continuous speech in CI users. Measures of neural tracking of the speech envelope reflect interesting properties of the listener’s perception of speech, such as speech intelligibility or attentional state. Successful decoding of neural envelope tracking will open new possibilities to investigate the neural mechanisms of speech perception with a CI.

## I. INTRODUCTION

The slow modulations of speech, also known as the speech envelope, are tracked by the neural oscillations in the listener’s brain (Aiken & Picton 2008, Peelle & Davis 2012). It is possible to reconstruct the speech envelope from recordings of neural responses to continuous speech, for instance using EEG or MEG (Ding & Simon 2012). This speech envelope decoding paradigm has been extensively investigated in recent years, as the decoded envelope may reveal information about the listener’s perception and processing of the stimulus. Some examples of factors that affect the neural tracking of the speech envelope are the stimulus intelligibility (Ding & Simon 2013, Vanthornhout et al. 2018), listener’s attentional state in a multi-speaker environment, (O’Sullivan et al. 2014), listener’s age (Presacco et al. 2016), and hearing loss (Petersen et al. 2016).

One population that would benefit greatly from an objective measure that reflects these factors are people with a hearing impairment. Besides possible diagnostic applications, another proposed application is the so-called neuro-steered hearing aid (Biesmans et al. 2017, Van Eyndhoven et al. 2017, O’Sullivan et al. 2017). Such devices automatically and autonomously adapt their audio signal processing settings based on, for instance, the user’s attention or speech understanding predicted from the neural recordings. With the rapid development of efficient distributed signal processing schemes (Bertrand 2015) and miniaturized EEG modules that can be integrated with hearing aids for recording of around-the-ear (Debener et al. 2015) or in-the-ear (Looney et al. 2012) EEG, the realization of such hearing aids becomes possible in the near future.

The same ideas can be applied to a cochlear implant (CI), which enables hearing in deaf or severely hearing impaired users through direct electrical stimulation of the auditory nerve. In this case, even the implanted electrodes could be used to unobtrusively monitor neural responses (Mc Laughlin et al. 2012). Additionally, an envelope tracking measure seems very appropriate for studying speech reception in CI users, since the audio envelope is the main speech feature that gets encoded in the electrical stimulation sequence of the implant (Wouters et al. 2015).

Nevertheless, no studies investigating speech envelope tracking in CI users have been presented so far. The major challenge to overcome when using EEG in CI users is the interference of electrical stimulation artifacts. Every stimulation pulse delivered by the implant causes a large artifact in the recordings. For transient evoked responses, techniques to deal with these artifacts exist. If very short stimuli (e.g. clicks) are used, the artifact decays before the response occurs, and no artifact removal is required. For longer stimuli such as tone bursts, speech utterances or natural sounds, the artifact (partly) overlaps with the evoked response and hinders the analysis. Several techniques to remove such artifacts have been proposed, such as template-based methods, which first estimate the artifact and then subtract it from the signals, (Friesen & Picton 2010, Mc Laughlin et al. 2012, 2013, Deprez et al. 2017*b*), and multi-channel technniques such as principal component analysis (Martin 2007) or independent component analysis (Gilley et al. 2006, Martin 2007, Sandmann et al. 2009, Viola et al. 2011).

When investigating steady state or stimulus tracking responses, electrical stimulation is continuously ongoing, causing the artifacts to obscure the entire EEG recording. To make matters worse, modulations in the artifacts directly correspond to modulations in the presented stimulus and are highly correlated with the neural responses of interest. This makes disentangling the artifact from the response difficult using the aforementioned techniques. Nevertheless, it is especially important to eliminate stimulus artifacts, or they will result in artifact-dominated responses that resemble the expected neural responses but are in fact false positives. In the case of electrically evoked auditory steady state responses (EASSRs), reliable responses can be obtained using a linear interpolation technique to remove the artifacts (Hofmann & Wouters 2010, Gransier et al. 2016, Deprez et al. 2017a). However, this technique has only been applied for stimulus pulse rates of maximally 500 Hz to 900 Hz, because the time interval between successive stimulation pulses should be long enough for the artifact to decay. This prevents the interpolation method from being applied to recordings using speech stimuli, as the pulse rates required for decent speech understanding are often an order of magnitude larger (e.g. 7200 Hz). Also other CI artifact removal techniques that work well for periodic EASSR stimulation such as Kalman filtering (Luke & Wouters 2017) are not easy to adapt for complex, high-rate speech stimulation.

In the present study, we demonstrate for the first time an EEG-based method for assessing neural tracking of the speech envelope in CI users. The novelty of our approach is in the elimination of the aforementioned CI artifacts, which is essential to obtaining a truly neural measure. The basic principle of our approach is to modify the electrical stimulus sequence such that short time segments without any stimulation appear, during which the artifact-free EEG can be sampled. Using this low-rate EEG, a conventional linear decoder is used to reconstruct the stimulus envelope and obtain a measure of neural envelope tracking. We also validate that the obtained EEG is free of stimulus artifact.

## II. METHODS

### A. Overview

The envelope decoding paradigm employed in the present study is illustrated in Fig. 1. In essence, two envelopes are obtained and compared to each other: one envelope reconstructed from EEG recordings, and one envelope extracted from the presented stimulus. The reconstructed envelope will resemble the stimulus envelope if it is encoded properly in the listener’s brain. As such, the correlation between reconstructed and real stimulus envelope is used as a measure of neural tracking of the speech envelope.

**Fig. 1.**
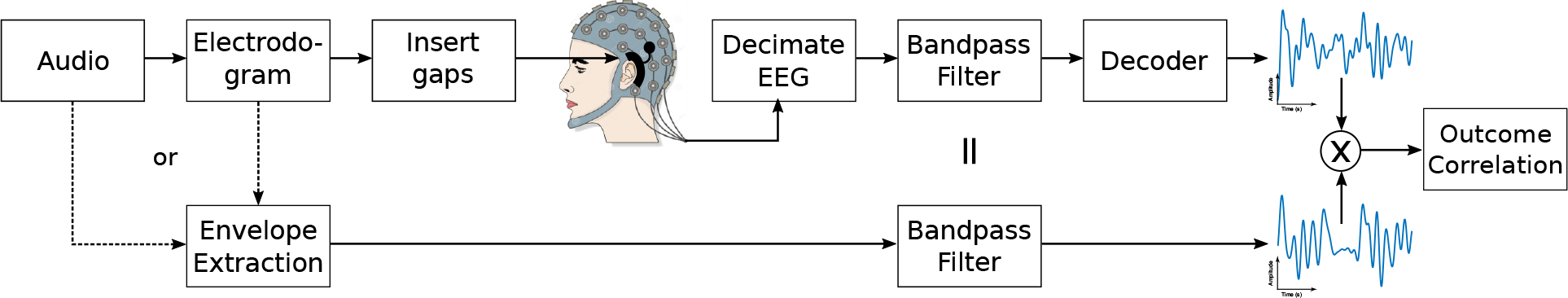
Schematic overview of signal processing pipeline.

Various methods to extract the stimulus envelope from an acoustic waveform exist (Biesmans et al. 2017). However, for CI users, those established methods may not represent the stimulus envelope very well, as the audio passes through many non-linear processing stages in the cochlear implant sound processor. The results using an acoustic-based stimulus envelope will be compared with results obtained using an envelope derived from the electrical pulse sequence.

The reconstructed envelope is obtained by applying a linear decoder to the EEG recordings. The decoder is trained on a training set consisting of EEG and the concurrently presented stimulus envelope, such that it “learns” the spatiotemporal relation between EEG and stimulus. The decoder can then be applied to a testing set of EEG in order to reconstruct the stimulus envelope. This reconstruction is then compared to the stimulus presented during the testing EEG.

It is not possible to directly use EEG data recorded in CI users for training and testing of the decoder due to the large artifacts originating from the electrical stimulation. If these artifacts are (partly) left in the EEG, the decoder is able to compute an accurate envelope reconstruction based on the artifacts, instead of based on the neural responses. In order to avoid such false positive results, temporal gaps were inserted in the electrical stimulus sequence. The EEG sampled within these gaps is free of stimulus artifact, and can be used for envelope decoding purposes. The trade-off between obtaining enough artifact-free EEG samples and not significantly degrading speech intelligibility by inserting these gaps was carefully balanced.

### B. Subjects

Eleven CI users participated in the present study. All subjects were native Flemish-speaking adults with at least 6 months of experience of listening with their cochlear implant. All subjects had implants from Cochlear Ltd.. Relevant details for all subjects can be found in Table I. The study was approved by the Medical Ethics Committee UZ / KU Leuven with reference number S57102.

**TABLE I.** SUBJECT DETAILS

| Subject | Age (years) | Gender | CI experience (years) | CI side | CI device |
| --- | --- | --- | --- | --- | --- |
| S1 | 46 | F | 3 | R | CI422 |
| S2 | 56 | F | 0.5 | R | CI522 |
| S3 | 58 | M | 2 | L | CI522 |
| S4 | 70 | M | 4 | L | CI24RE (CA) |
| S5 | 46 | F | 5.5 | R | CI24RE (CA) |
| S6 | 21 | M | 16 | R | CI24M |
| S7 | 20 | M | 18 | R | CI24M |
| S8 | 61 | F | 1 | L | CI522 |
| S9 | 73 | F | 1 | L | CI522 |
| S10 | 76 | M | 1.5 | L | CI522 |
| S11 | 58 | F | 1 | R | CI522 |

### C. Stimuli

The speech stimuli were stories^1^ narrated by a female Flemish speaker. The Nucleus Matlab Toolbox (NMT) (Swanson & Mauch 2006) was used to convert the audio files into electrical stimulation sequences similar to a real CI sound processor. Briefly, a CI sound processor splits the input audio waveform into frequency subbands, which are mapped to 22 electrode channels. In each of these channels, an electrical pulse train is modulated with the envelope of its allocated frequency subband. As such, this scheme mimics the frequency-selective sound processing of the cochlea. The output of the sound processor is a modulated pulse sequence for every electrode channel, known as an electrodogram. An example electrodogram is shown in Fig. 2(a).

**Fig. 2.**
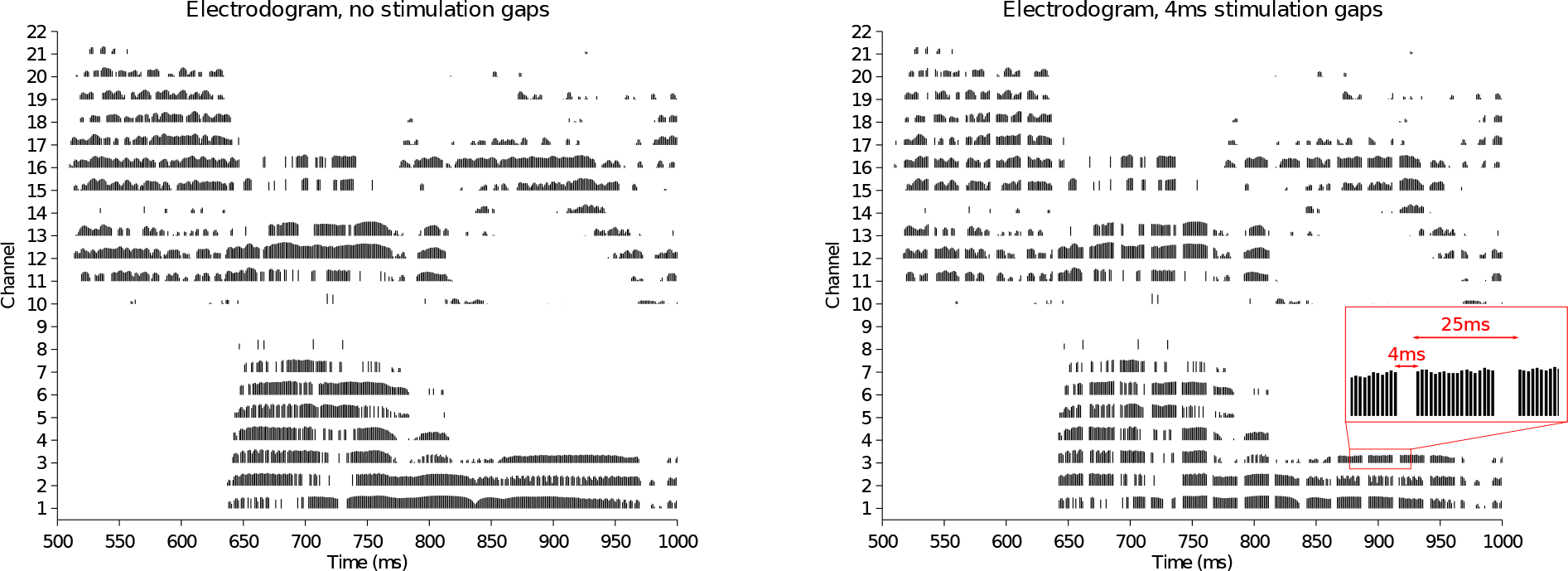
Illustration of gap insertion in the electrodogram. (a) Electrodogram after processing half a second of speech audio. Each channel of the electrodogram roughly corresponds to a pulse train modulated with the subband envelope assigned to that channel. As such, each channel conveys frequency-specific information to the different frequency-sensitive areas of the cochlea. (b) Temporal gaps are inserted in the stimulus sequence by omitting the pulses across all channels for a few milliseconds.

The subjects’ clinical MAPs were used to set all sound processing parameters. The MAP of a CI user is a collection of subject-specific implant settings required for the conversion of input sound to electrode stimulation. The most important settings are the Threshold (T) and Comfort (C) levels for each electrode, expressed in current units (cu). The T-level of an electrode is the lowest stimulation level that results in a perceivable sound. The C-level of an electrode is the highest stimulation level that results in a comfortably loud sound. Other parameters are number of active electrodes, frequency allocation per electrode, stimulus pulse width, pulse rate, etc. The intensity of the input audio was adjusted such that the resulting electrical stimulus corresponded to an acoustic signal of 60 dB SPL at the CI microphone.

The experiments were conducted using the open source APEX software (Francart et al. 2008), which allows interfacing with Nucleus devices through the Nucleus Implant Communicator (NIC). The stimulation sequences were presented to the recipient’s CI using a research speech processor (L34) provided by Cochlear Ltd.. The stories were 15 minutes long and were presented in blocks of 5 minutes. After every block, the subject was questioned about the content of the story to promote the subject’s attention.

### D. Recording

All recordings were performed in an electrically shielded Faraday cage using a BioSemi ActiveTwo amplifier with 64 electrodes. Data was recorded at a sample rate of 16 384 Hz, except for subject S1 whose recordings were obtained at 8192 Hz. A trigger signal emitted by the stimulating device was recorded along with the EEG signal for synchronization. Because EEG preprocessing operations such as referencing and filtering can smear out the CI stimulation artifact both spatially and temporally, all EEG preprocessing was performed after eliminating the CI stimulation artifacts (see section II-E). There were multiple electrodes for each subject that did not make a good contact with the scalp due to the coil and earpiece of the CI. The affected electrodes were different for every subject. To avoid manual picking of good and bad electrodes, only the 32 electrodes on the side contralateral to the stimulated ear were used in the analysis.

### E. Elimination of cochlear implant artifacts

#### 1) Problem description

The CI applies electrical stimulation pulses on every electrode, with pulse timing and magnitude specified by the pulse sequence generated from the audio signal (see Fig. 2(a)). Every individual pulse causes an artifact in the recorded EEG. While the applied pulses are rectangular and symmetrically biphasic, the morphology of the recorded artifacts is altered by the finite recording bandwidth, nonlinearities in the EEG amplifier, and effects of the subject’s head such as volume conduction. While the phases of the stimulated pulses are typically only 25 µs or 37 µs long, the artifacts in the EEG recording are smeared out over several milliseconds. Because of the high total pulse rates (e.g. 7200 Hz) required for decent speech understanding, the artifacts largely overlap with each other.

Fig. 3(c) illustrates the manifestation of CI artifacts in a single EEG channel. The corresponding acoustical speech signal and its envelope are shown in resp. Fig. 3(a) Fig. 3(b). Both the stimulus artifact and the targeted envelope-tracking neural oscillations correlate strongly with the stimulus envelope. Without CI artifact removal, it is a trivial task to reconstruct the envelope from the EEG, as the linear decoder would just use the artifacts. It is crucial to eliminate the artifacts to obtain a measure that reflects neural responses to the speech envelope instead of an artifact-dominated reconstructed envelope.

**Fig. 3.**
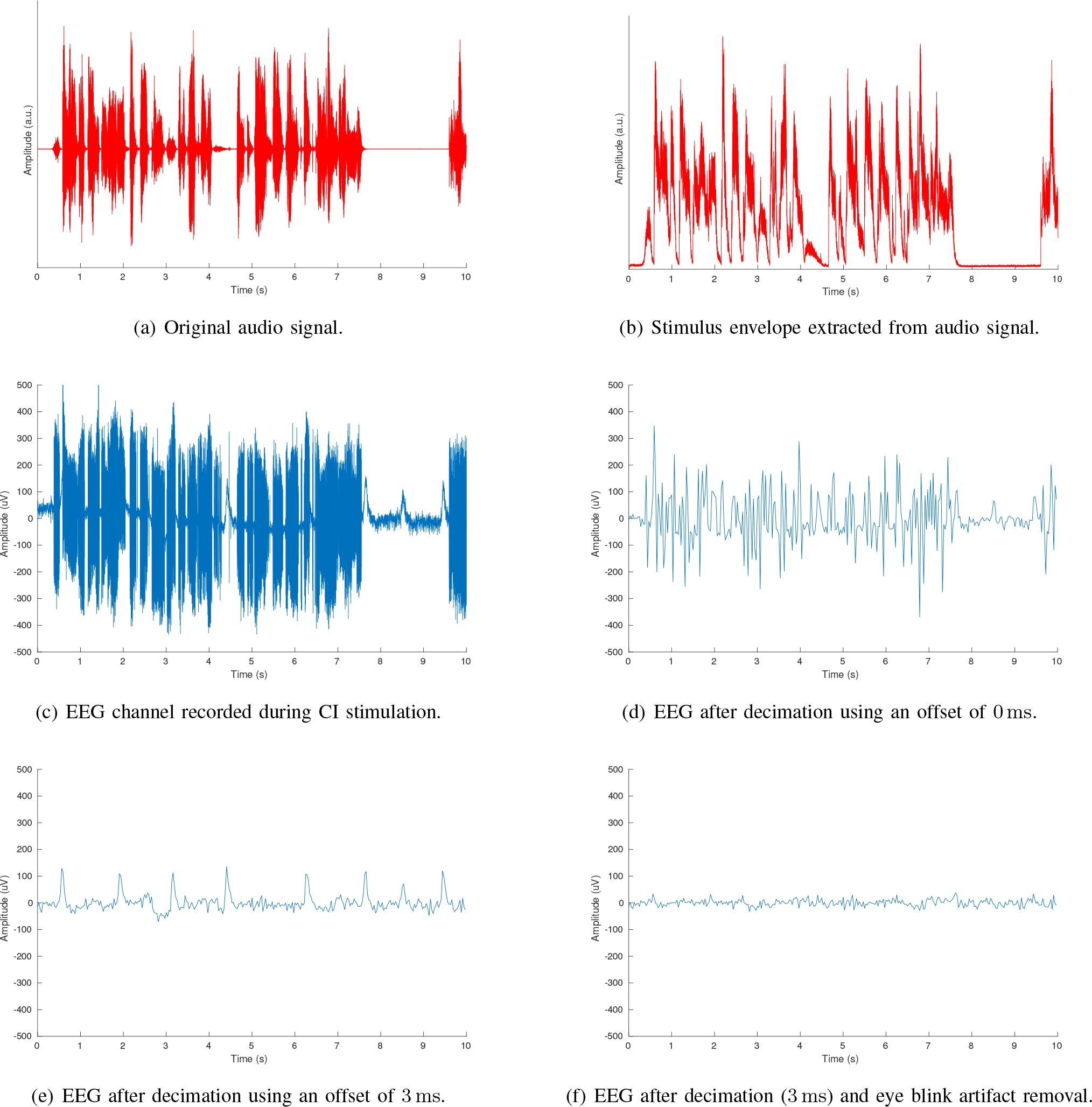
Examples of audio and EEG signals. (a) Original audio signal. (b) Stimulus envelope extracted from audio signal. (c) Single EEG channel signal (Fpl) recorded during CI stimulation. The large artifacts are resemble the audio stimulus and stimulus envelope. There are also eye blink artifacts in addition to the CI stimulus artifacts. (d) The same EEG signal after decimating using an offset of 0 ms. Because the offset is too small, there is residual stimulus artifact in the EEG. (e) The same EEG signal after decimating using an offset of 3 ms. This offset is sufficient so that the CI artifact has decayed. (f) Decimated EEG signal (3 ms) after eye blink artifact removal. This artifact-free EEG will be bandpass filtered and used for the decoder.

#### 2) CI artifact removal strategy

The strategy to obtain EEG free of CI artifacts consists of two parts. Firstly, temporal gaps are inserted in the stimulation sequence by periodically interrupting the stimulation for a few milliseconds. In practice, this was done by omitting stimulus pulses across all channels of the electrodogram, as illustrated in Fig. 2(b). The omission of these pulses is characterized by the duration of each gap (e.g. 4 ms) and the frequency at which the gaps occur (e.g. 40 Hz means a gap is inserted every 25 ms).

Secondly, the recorded EEG is decimated by retaining only samples within the stimulus gaps. Because there is no stimulation during these gaps, this should result in EEG signals free of CI artifacts. In practice, some time margin should be respected to allow the artifacts to decay after the stimulation is interrupted. The decimation process is illustrated in Fig. 4. The raw EEG signal is severely affected by CI artifacts, but drops back to normal EEG amplitudes during the stimulus gaps. The dotted vertical lines represent the decimation sampling moments, i.e., the EEG samples coinciding with these lines are retained while all other samples are discarded. These sampling moments should fall within the gap, but their offset from the gap starts can be chosen freely. The inset figures in Fig. 4 show that there is a transient artifact decay at the beginning of the gap. Therefore, it is important to choose the sampling moments near the end of the gap, such that the transients have decayed and do not affect the retained samples. This effect is illustrated in Fig. 3: the raw EEG signal with artifacts from in Fig. 3(c) is decimated by retaining a single sample in every gap. When the samples are taken at the start of the gap (i.e. an offset of 0 ms relative to gap start), residual artifacts remain in the decimated EEG (Fig. 3(d)). When selecting a choosing a larger offset (e.g. 3 ms), the artifact has decayed at the moment of sampling and does not appear in the decimated EEG (Fig. 3(e)). Note that the decimation eliminates CI artifacts, but other artifacts such as eye blinks still have to be removed in a later stage to obtain a clean EEG signal as shown in Fig. 3(f) (see section 11-F).

**Fig. 4.**
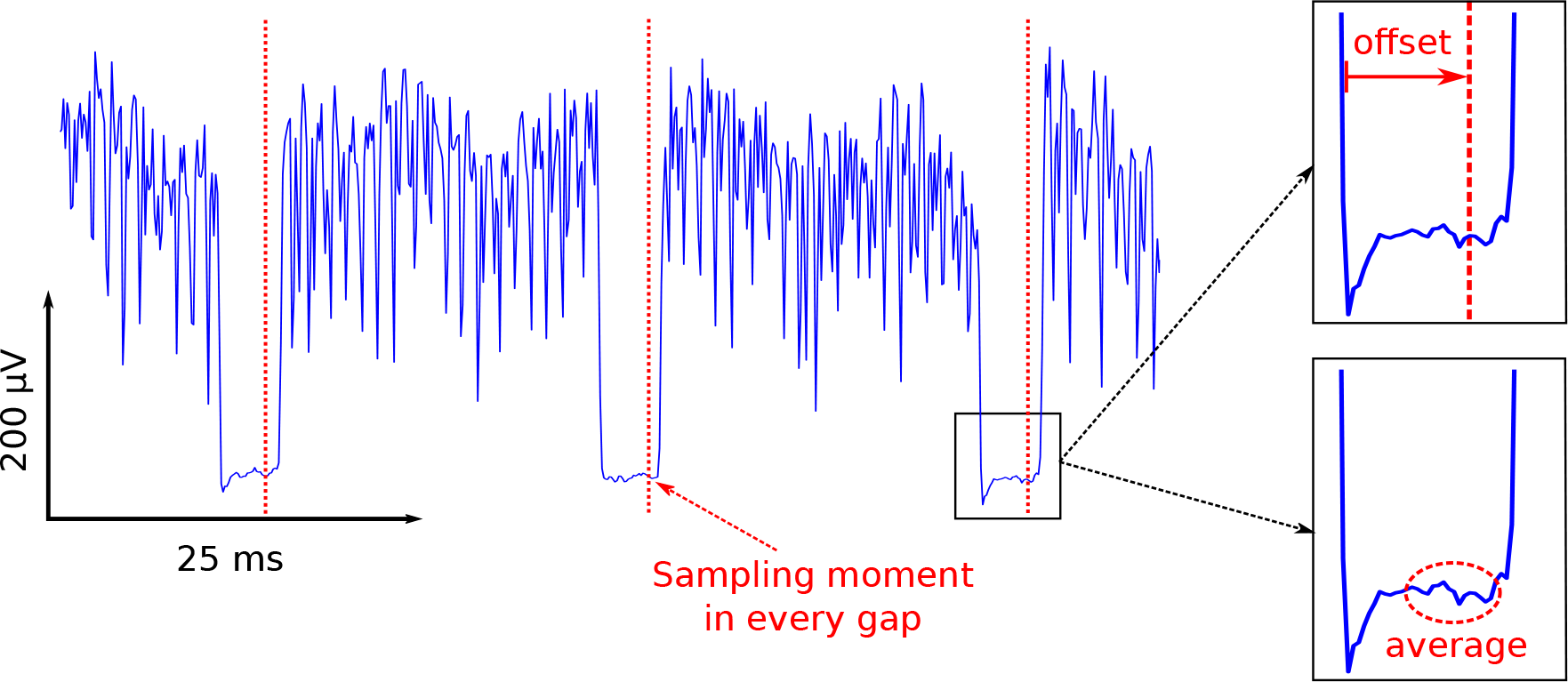
Illustration of the EEG decimation process. In this example, a gap frequency of 40Hz is used, such that stimulation is interrupted every 25Hz. In each gap, an EEG sample is chosen to retain, resulting in an EEG sample rate of 40Hz. The sample to be picked in each gap can be chosen in two ways, as shown in the inset figures: either a single sample at a certain offset from the gap start (top), or an average of multiple samples (bottom).

A slightly different method is to take an average of multiple samples in each gap and use these samples as the decimated EEG. In the sequel, this condition will be referred to as the "average" decimation condition. The samples in the last millisecond of the gap are averaged together to avoid the transients at the start of the gap. Because of the averaging, the noise inherent to EEG will be reduced, resulting in a better estimate of the neural response to the speech envelope.

#### 3) Gap parameters and effect on speech intelligibility

The choice of the gap duration and frequency affects the resulting EEG after decimation. The gap frequency determines the EEG sample rate after decimation, as one sample per gap is retained. Because the targeted envelope-tracking responses are present in the lowest EEG frequencies (e.g. up to 10Hz), the required minimal EEG sample rate after decimation is 20Hz due to Nyquist’s sampling theorem.

The duration of the gaps needs to be sufficient to obtain artifact-free EEG samples, i.e. the artifacts caused by the stimulus pulses right before the gap should have decayed when the samples in the gap are picked. This decay follows an exponential time course and is influenced by several factors such as stimulus intensity, EEG sample rate, and subject-related factors such as tissue impedances around the stimulating electrodes. As artifacts were expected to decay within 1 ms to 2 ms post-stimulus (Deprez et al. 2017a, Hofmann & Wouters 2010) and including some margin for error, gaps lengths between 2 ms to 5 ms were tested.

Evidently, the gap duration and frequency cannot be increased too much as the intelligibility of the speech stimulus would decline. Due to the pulsatile nature of the stimulation, the speech intelligibility of CI users is quite robust against omission of some stimulus pulses. Nevertheless, the effect on speech intelligibility was assessed behaviorally for each subject before starting the EEG recordings. The gap parameters where gradually increased until the subject reported that the stimulus either became more difficult to understand, or became uncomfortable to listen to. Next, **it** was verified that speech intelligibility was not negatively affected by inserting gaps with the selected parameters. This was done by comparing sentence scores for a standardized speech material (LIST sentences, (Van Wieringen & Wouters 2008)) presented with and without gaps.

#### 4) Validation of artifact removal

The removal of CI artifacts is validated in two ways. Firstly, the envelope decoding scheme is repeated for different sampling moments during the decimation step. In other words, the sampling offsets as in Fig. 4 are varied from 0 ms up to the gap length. As the offset increases, the contribution of CI artifact in the sample at that offset will decrease, while the contribution of the neural response remains constant. By observing the outcome measure over a range of offsets, the offset after which the stimulus artifacts have decayed and do not affect the response anymore can be determined.

Secondly, additional EEG recordings are made while the stimulus was presented below the subject’s threshold level, which will be referred to in the sequel as sub-T EEG. Like the other recordings, the sub-T EEG contains CI artifacts which are highly correlated with the stimulus envelope. However, because the stimulus was inaudible to the subject, the sub-T EEG can not contain any neural responses to the stimulus. If any envelope-tracking response is observed in the sub-T EEG, **it** must be a false positive response caused by the CI artifacts.

### F. Signal processing

The neural tracking of the speech envelope is measured by correlating the stimulus envelope with the envelope reconstructed from the EEG recordings. Fig. 1 shows the signal processing steps used to obtain both of these envelopes. All signal processing was implemented in MATLAB.

The stimulus envelope was derived in two different ways: either based on the acoustic stimulus waveform as in the literature, or based on the stimulus electrodogram. In the sequel, we will refer to these extracted envelopes as the acoustic and electrical envelope, respectively. For the acoustic envelope extraction, a gammatone filterbank followed by a power law with exponent 0.6 was used, based on the comparison between speech envelope extraction methods for envelope decoding in (Biesmans et al. 2017). The gammatone filterbank implementation from the Auditory Modeling Toolbox (AMT) (Søndergaard & Majdak 2013) was used. The electrical envelope was hypothesized to be a more realistic envelope representation for CI stimulation as it is derived from the electrodogram, which takes into account the non-linear processing from the CI sound processor. The electrical envelope is obtained by adding together the modulated pulse sequences of all channels of the electrodogram. The obtained stimulus envelope (either acoustic or electrical) is downsampled to the EEG sample rate and filtered with a bandpass filter with a passband encompassing the EEG delta band (0.5 Hz to 4 Hz), which is important for speech processing.

The reconstructed envelope is derived from the EEG recordings. Firstly, CI artifacts are removed by decimation as described in section II-E. A multi-channel Wiener filter-based algorithm (Somers et al. 2018) is applied to remove eye blink artifacts, which are low-frequency artifacts with significant spectral power in the frequency band of interest. Next, the EEG is filtered with the same bandpass filter used for the stimulus envelope. The low-rate, bandpass filtered EEG and envelope are jointly used to train a linear decoder (see section II-G), which is used to obtain the reconstructed envelope.

While the electrical envelope already takes into account some of the non-linear processing in the CI sound processor, more non-linearities are introduced while encoding the envelope in the brain. The envelope reconstruction may be improved by adding an additional non-linear operation to the reference envelope extracted from the stimulus. While the exact non-linear operation of the brain is unknown, a simple non-linear model to include in the electrical envelope extraction is a power law, similar to the extraction of the acoustic envelope. The effect of applying a power law to the electrical envelope to improve decoding results will be investigated for a range of different exponents.

Finally, a bootstrapped Spearman correlation is computed between real and reconstructed stimulus envelopes. Bootstrapping is performed by Monte Carlo resampling of both envelopes (*N* = 100). The median of the boot-strapped correlations is taken as a measure of the strength of neural tracking of the speech envelope.

### G. Training and validation of decoder

The linear decoder is a spatiotemporal filter that combines all of the *M* EEG channels and their time-delayed versions together into a single estimate of the stimulus envelope, i.e.

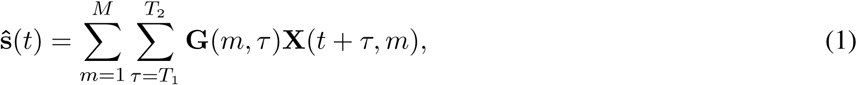

where **X** (*t, m*) is the EEG sample in channel *m* at time *t*, **G** is the linear decoder matrix, and s is the estimate of the stimulus envelope **s**. The time window [*T*_1_, *T*_2_] determines all time delays to be included in the decoder and is referred to as the integration window. Commonly, *T*_1_ is chosen *≥* 0 ms to ensure causality and *T*_2_ is chosen to be several tens or hundreds of milliseconds to take into account the expected duration of neural processing.

The decoder matrix **G** can be found by minimizing the least squares error between actual and estimated stimulus envelopes, i.e. solving

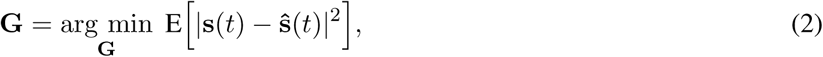

where E[●] denotes the expected value operator. The solution to this minimization problem corresponds to the least squares fit of **X***_τ_* to **s**, which can be practically calculated as

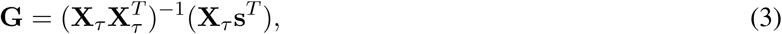

in which **X***_τ_* is the matrix containing the EEG recordings with all possible delays *τ* in the integration window stacked together, i.e.

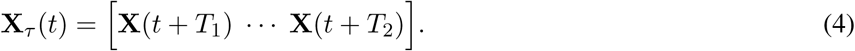

Zero-padding is applied to ensure all stacked time-delayed EEG recordings have the same length. All decoders are calculated using the mTRF toolbox (Crosse et al. 2016). The integration window is set from 0 ms to 250 ms.

Decoders are trained and tested using a leave-one-out cross-validation scheme illustrated in Fig. 5. The 15 minutes of EEG recording and stimulus envelope are split into two mutually exclusive sets of training and testing data. The training data is 13 minutes long and is used to train a linear decoder according to (3). The obtained decoder is applied to the 2 minutes of testing set EEG to reconstruct the envelope according to (1). The reconstructed envelope is then correlated with the testing set stimulus envelope to obtain the measure of neural tracking. The analysis is repeated for different partitions of training and testing sets, such that the testing set does not overlap with one previously used. As the story has a duration of 15 minutes, 7 non-overlapping testing sets of 2 minutes can be used, resulting in 7 trials with associated decoders and outcome correlation measures.

**Fig. 5.**
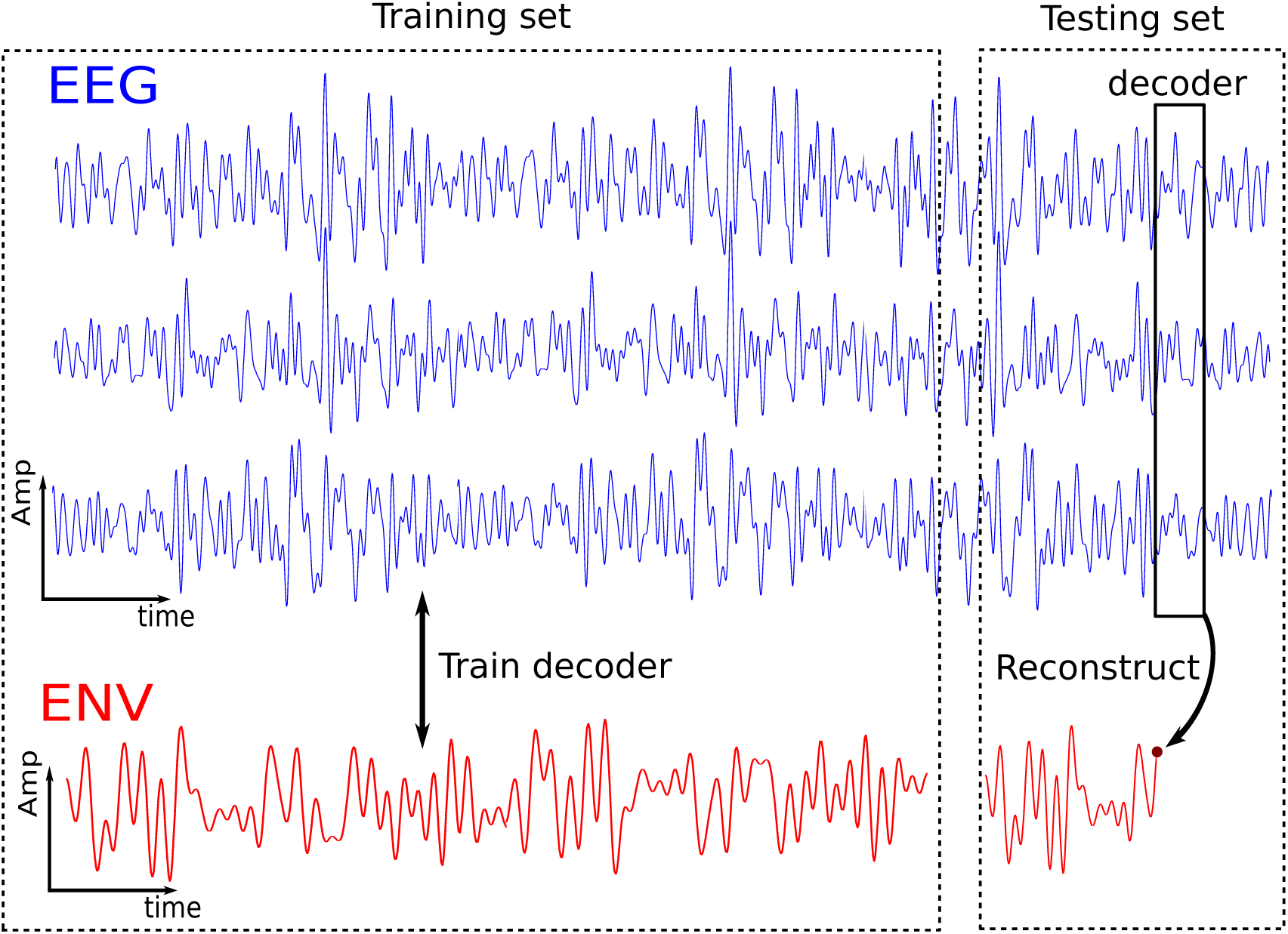
Illustration of training and testing scheme of the linear decoder. Note that only three EEG channels are depicted, and their time-delayed versions are omitted from the illustration. The EEG and the concurrently presented envelope (ENV) are split in a training set (13 min) and a testing set (2 min). The training set is used to "teach" the decoder the stimulus-response relationship. This decoder is then applied to the testing set to reconstruct the envelope. The scheme is repeated multiple times for different training and testing set segmentations.

Each decoder obtained using the leave-one-out cross-validation is also applied to the sub-T EEG, resulting in 7 correlation values for the sub-threshold stimulation condition. To determine if the obtained correlations are significant (using α = 0.05), significance levels are obtained by correlating random permutations of the real and reconstructed envelopes 1000 times and taking the 2.5^th^ and 97.5^th^ percentiles. Correlations between these two percentiles are not significant, which should be the case for correlations obtained using the sub-T EEG, because this stimulation was inaudible for the subjects. A sub-T correlation above the significance level indicates that CI artifact were not removed successfully and caused a false positive response.

## III. RESULTS

### A. Perceptual effect of small gaps in the stimulus sequence

The insertion of gaps in the stimulus as in Fig. 2 is required to sample artifact-free EEG, but should not reduce speech intelligibility too much, as this would also negatively affect the neural tracking measure. The effect of the gaps was assessed behaviorally. As longer and more frequent gaps were inserted, subjects reported that the speaker’s voice sounded increasingly artificial or robotic. The gap length and frequency were increased until the subject indicated that the stimulus became less intelligible or unpleasant to listen to. The resulting gap parameters were similar across subjects: a gap length of 4 ms was found for all subjects, and a gap frequency of 40 Hz was found for all but two subjects (S5 and S6 had a maximal comfortable gap frequency of 32 Hz).

The effect on speech intelligibility was measured using standardized speech sentences for 8 out of 11 subjects (all except S1, S6 and S7). Using unaltered sentences, the subjects achieved speech intelligibility scores of 96.5 ± 4.6% (mean ± standard deviation). For sentences with gaps, the scores were 97.7 ± 2.7%. There was no significant difference between the scores (Wilcoxon signed rank test, *p* = 0.82). The mean difference between scores of unaltered and altered stimuli is –1.25 (95% confidence interval = [–6.0, 3.0]).

### B. Effect of stimulus artifacts on neural tracking measure

Fig. 6 shows the outcome correlation measures obtained for a representative subject, as a function of sampling offset used for the decimation. For each offset, there are seven correlations for the above-threshold condition and seven for the sub-threshold condition, resulting from the leave-one-out cross-validation procedure as described in section 11-G. The significance levels are indicated with the horizontal dotted lines. For this subject, the gaps had a duration of 4 ms and a frequency of 40 Hz, and the electrical envelope was used.

**Fig. 6.**
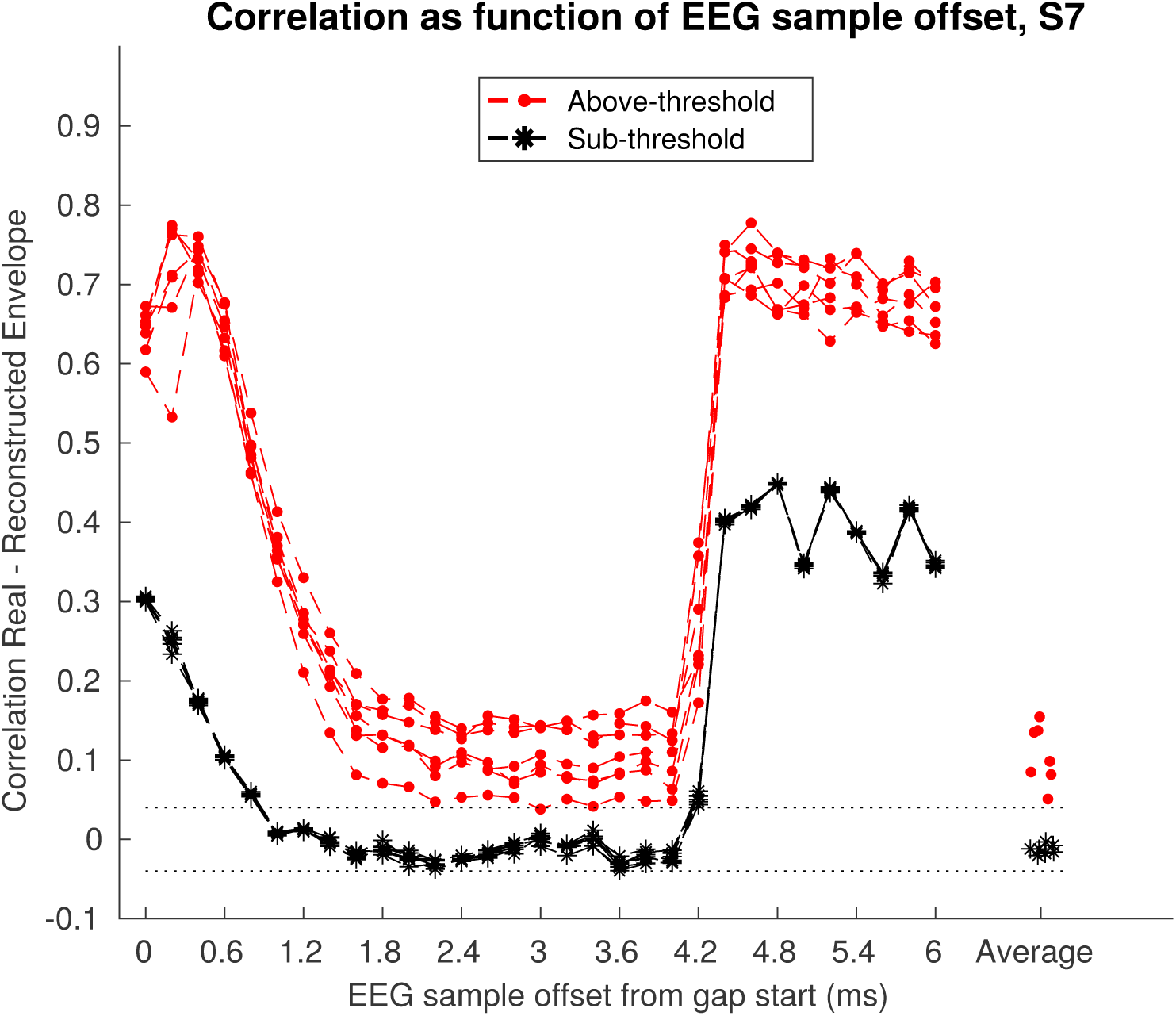
Outcome correlation measure as function of sampling offset during decimation step for representative subject. Sampling offset is varied from 0 ms to 6 ms. The average decimation condition is shown as well on the right.

Both above-threshold (i.e. using the subject’s everyday, clinical settings) and sub-threshold correlations decline for small offsets (0 ms to 2 ms), resembling an exponential decay as expected. In this range, the decimated EEG still contains residual stimulus artifacts and the correlations are artifact-dominated and thus too high. For larger offsets, the outcome correlations flatten out, i.e. they become invariant to the sampling offset, indicating that the CI artifact has decayed. The correlations obtained with the regular above-threshold stimulation are above the significance level, whereas those using below-threshold stimulation are not. After 4 ms stimulation resumes, which causes the correlations to return to inflated, artifact-dominated values similar to those at the start of the gap at offset 0 ms. Results for the average decimation condition are shown as well and are similar to results obtained for offsets between 3 ms to 4 ms.

Fig. 7 shows a similar analysis on group level. Fig. 7(a) and 7(b) show results obtained using the electrical envelope and the acoustic envelope, respectively. The mean and standard deviation of the outcome correlation measures across all subjects are shown as a function of sampling offset. Results for offsets up to 4 ms are shown, as this is the gap length for all subjects, as well as subject-specific results for the average condition. The group level results show a similar course as the subject-specific result in Fig. 6. After an initial artifact decay, the correlations flatten out. Above-threshold stimulation results in correlations above the significance level, while sub-threshold stimulation does not result in meaningful correlations.

**Fig. 7.**
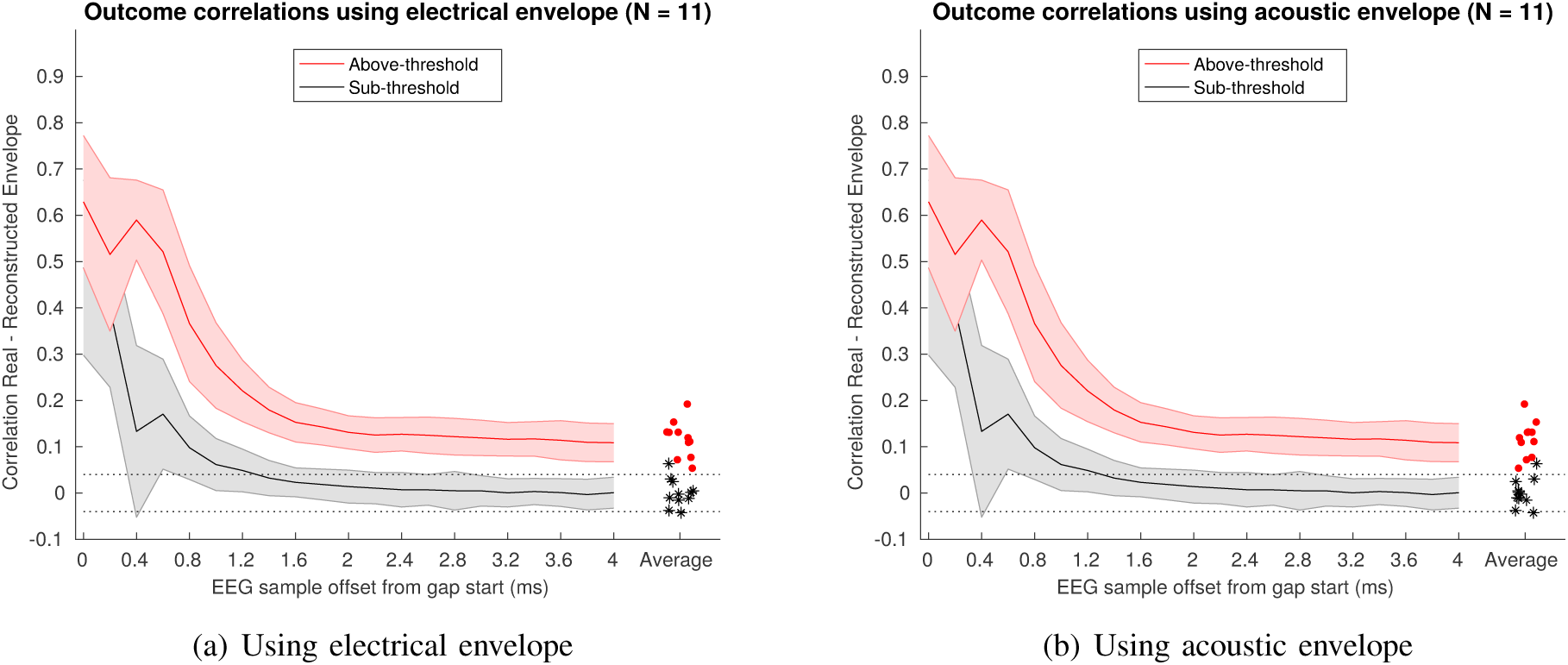
Group level (N = ll) correlations as function of sampling offset during decimation. Solid lines represent the mean across subjects, shaded area widths represent the standard deviation across subjects. Additionally, individual results for the average condition are shown and are jittered for increased visibility. (a) Using electrical envelope derived from electrodogram. (b) Using acoustic envelope derived from original audio waveform.

### C. Choice of reference stimulus envelope

By comparing the results in Fig. 7(a) and 7(b), it can be observed that after successful CI artifact removal (i.e. when the exponential has decayed), the average correlations are not affected too much by choice of envelope. On the other hand, when there is still residual artifact in the EEG, the correlations tend to be slightly larger when using the electrical envelope.

The effect of applying different a power law with varying exponents to the electrical envelope is shown in Fig. 8. The median outcome correlation measure for each subject, using the average decimation condition, is shown as a function of exponent applied to the reference electrical envelope. A power law exponent of 1 does not alter the envelope and corresponds to the average condition results shown in Fig. 7(a). Interestingly, the correlation measure reaches a maximum for most subjects when using a low exponent in the 0.2 - 0.5 range, but this is not true for all subjects. Also note that for subject S5, whose correlations were below the significance level using an exponent of 1, the results become significant using smaller exponents.

**Fig. 8.**
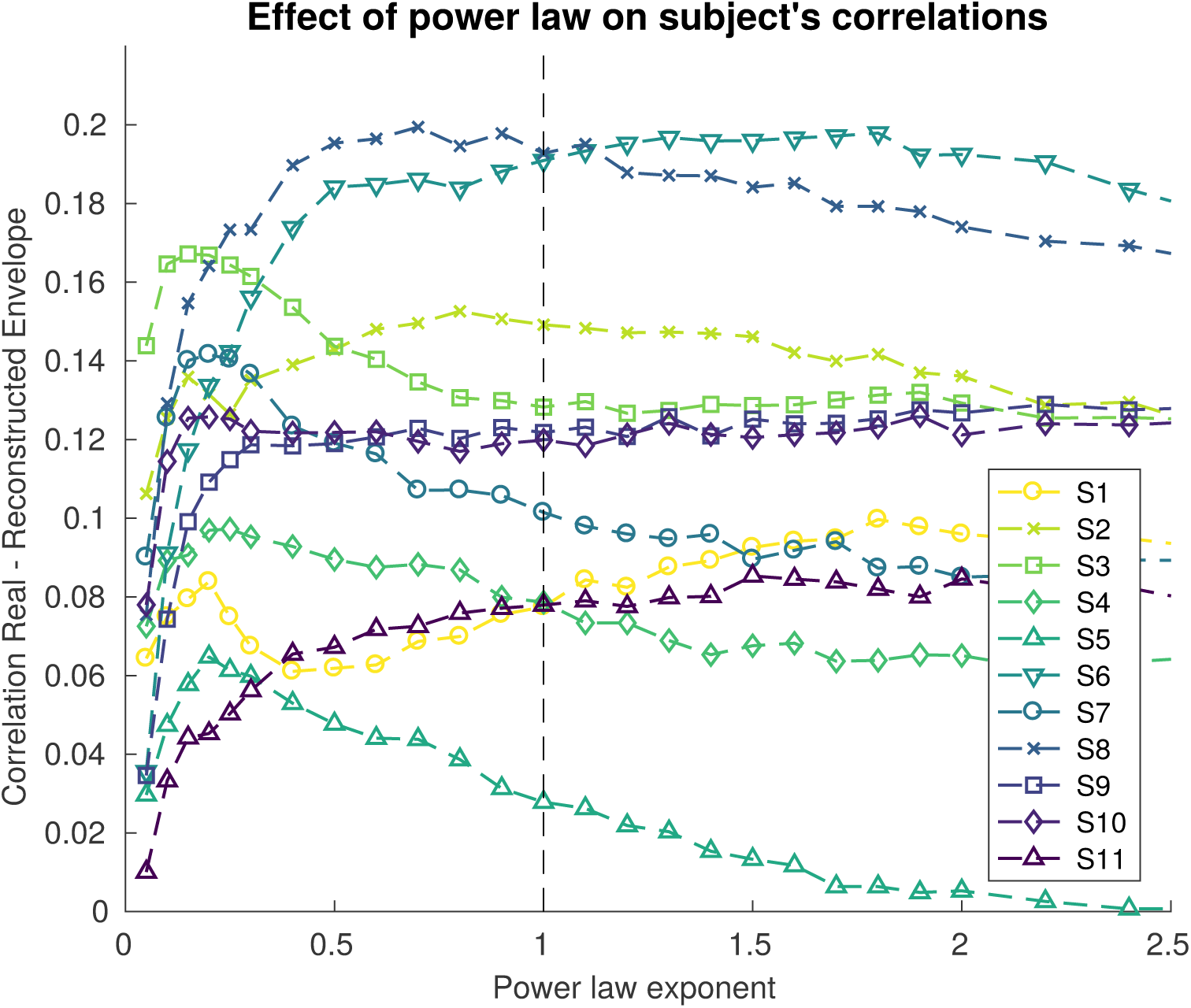
Effect of applying a power law to the electrical reference electrode. Each trace corresponds to the median correlation measure for a single subject using the average decoding condition. The vertical dashed line corresponds to the default exponent of 1, i.e. no power law applied to the envelope.

### D. Can a better choice of integration window suppress CI artifacts?

Due to the choice of decoder integration window, EEG samples with latencies of 0 ms up to 250 ms post-stimulus are used to reconstruct the speech envelope. As evidenced by Fig. 7, CI artifacts decay over a course of several milliseconds. On the other hand, neural processing of speech occurs at longer latencies in the integration window.

An intuitive strategy to suppress the influence of CI artifacts on the decoded responses would be to restrict the integration window to, for example, 25 ms to 250 ms. This would force the decoder to reconstruct the envelope based on longer-latency responses in the EEG, and prevents the instantaneous CI artifacts to contribute to the reconstructed envelope.

While appealing at first, this strategy fails to prevent CI artifacts from dominating envelope reconstruction. The autocorrelation of the speech envelope shows how well the envelope (but also the CI artifact) correlates with itself at certain latencies. If this autocorrelation still has a significant contribution at e.g 25 ms, then the CI artifacts at this latency have a predictive value for reconstructing the current sample of the speech envelope. As such, restricting the start of the integration window to 25 ms will not fully remove the influence of CI artifacts on the envelope reconstruction if the envelope autocorrelation is not zero at 25 ms.

The effect of manipulating the integration window is illustrated for one subject in Fig. 9. The outcome correlation is shown as a function of the start of the integration window used. The length of the integration window is kept constant at 250 ms, e.g. 25 ms on the horizontal axis means an integration window of 25 ms to 275 ms was used. This analysis is done for EEG with and without CI artifacts, i.e. decimated using a very small and a sufficiently large offset. respectively. Additionally, the autocorrelation of the band-pass filtered stimulus envelope is shown with a dashed line. In Fig. 9(a), the outcome correlations are dominated by CI artifact and they oscillate similarly to the stimulus autocorrelation as function of the integration window offset. This observed correspondence between the outcome correlation and the stimulus autocorrelation is not seen in the Fig. 9(b), where the CI artifact is removed.

**Fig. 9.**
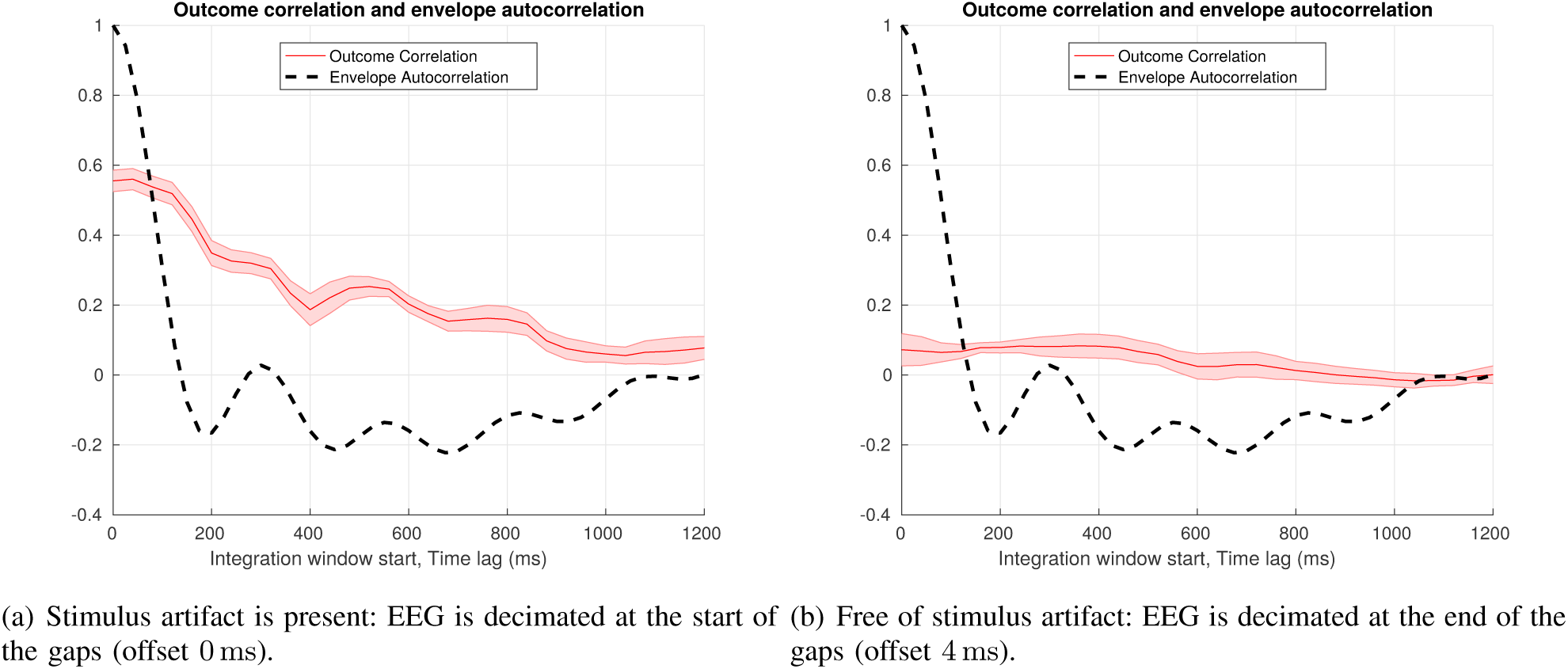
Envelope autocorrelation and outcome measures as function of the start of the integration window for EEG (a) containing CI artifacts and (b) free of CI artifacts.

## IV. DISCUSSION

A measure of neural tracking of the speech envelope in CI users was investigated. Stimulus artifacts were avoided by altering the electrical stimulus sequence to allow the sampling of short EEG segments free of artifacts. Furthermore, we investigated the effect of using different reference envelopes for training and testing the decoder, as it is not trivial which one better represents the encoded envelope after the non-linear processing in the subject’s CI and auditory pathway.

### A. Validation of the stimulus artifact removal

The CI stimulus artifacts were removed by periodically interrupting the electrical stimulation and sampling the EEG during those gaps. Intuitively, this leads to artifact-free EEG because there is no stimulation occurring when the EEG is sampled. However, due to the finite EEG recording bandwidth and nonlinearities during acquisition, it is possible that stimulus artifacts right before the gaps smear out and extend into them. To investigate these artifact tails, the sampling offset in the gap was varied (e.g. Fig. 6 and 7). The outcome measure shows an artifact-dominated tail that decays exponentially as the sampling offset increases. After a few milliseconds, the correlations plateau, indicating that the artifact has mostly decayed and the outcome measure is dominated by neural responses. While it is likely that the artifact is eliminated, this method cannot guarantee that the artifact is completely eliminated from the EEG samples, because theoretically an exponential decay continues on infinitely. However, these figures show that the contribution of the artifacts relative to the neural responses has become so small that it does not significantly alter the outcome measure anymore.

Similar conclusions can be drawn from the sub-threshold analysis. In these recordings, there is certainty that there is no neural response to the auditory stimulus, while there are stimulus artifacts. The outcome correlations again decay exponentially and plateau afterwards. The exponential decay starts at a lower amplitude, as the sub-threshold stimulation has lower intensity, but has a similar time constant, i.e. it plateaus at an offset similar to the above-threshold recordings. However, in this case the amplitudes of the plateau fall below the significance level. This proves that even if there would be any residual artifact, it is not strong enough to generate a significant false positive response.

Changing the decoder integration window to exploit the delay of neural auditory processing versus the instantaneous artifacts is not successful for removing artifacts (see Fig. 9). If there are any stimulus artifacts in the EEG, moving the start of the integration window to longer latencies still results in abnormally large outcome correlations, even when going past 500 ms where no important responses to the speech envelope are expected anymore. These false positive results may seem surprising given that the instantaneous artifacts cannot contribute to the envelope reconstruction, but they can be explained by considering the autocorrelation of the stimulus envelope (see section III-D). When using decimated EEG without stimulus artifacts (Fig. 9(b)), the outcome correlations have lower, more realistic values and do not seem to follow the oscillations in the stimulus autocorrelation.

### B. Choice of reference envelope

To obtain the measure of neural tracking of the speech envelope, a reference envelope is needed. This reference envelope is used both for training the decoder and to correlate the reconstructed envelope with. The reference envelope is extracted from the speech stimulus, but there are many possible ways of doing so. In studies with normal hearing participants, the envelope is derived from the acoustic speech signal. In CI users, the matter is complicated further since the acoustic stimulus is processed in multiple non-linear stages in the sound processor. This makes it difficult to model exactly how the stimulus envelope is transmitted to the auditory nerve and how it will be encoded in the neural oscillations.

The acoustic envelope was derived using a gammatone filterbank followed by a power law, which yielded the best results in (Biesmans et al. 2017). For deriving a reference envelope from the electrical stimulus sequence, no methods are described yet in literature. We opted for the simplest approach by adding together the different channels of the electrodogram. This already takes into account some non-linear processing by CI, but it does not take into account the subject-specific mapping of each channel level to the different stimulation electrodes. Comparing outcome correlations obtained using the acoustic or the simple electrical envelopes, there are not apparent differences when the CI artifact has been eliminated (see Fig. 7). If there is still CI artifact left in the EEG (for small offsets), the correlations are slightly larger for the electrical envelope. This is logical, as the electrical envelope is a better representation of the stimulus artifacts.

In an attempt to improve decoding results using the electrical envelope, a power law with varying exponent was applied. This can substantially improve the decoding, but the optimal exponent differs between subjects (see Fig. 8). The results for the majority of the subjects benefit most from smaller exponents in the 0.2-0.5 range, but this is not true for all subjects. It is clear, however, that using a power law and selecting a subject-specific exponent can greatly improve the outcome correlations. We can speculate that this inter-subject variability relates to differences in perception of the stimulus and its encoding in the brain. For instance, it is known that CI users can have different loudness growth functions. The power law was introduced as a simple model to incorporate some non-linear brain processing into the reference envelope. Note that if the neural processing of the speech envelope would be strictly linear, the power law would be obsolete since the linear decoder can model linear processes.

### C. Limitations and implications of the presented study

There are some limitations to the demonstrated method for eliminating the stimulus artifacts. Firstly, the stimulus needs to be adapted to include gaps. While it was verified that the subjects’ speech intelligibility did not deteriorate, the alterations make the speaker’s voice sound more artificial (i.e. “robot-like”) and potentially increase listening effort. For experimental purposes this is not an issue, but for possible clinical or long-term monitoring applications, audible stimulus manipulation is undesirable. Secondly, the quality of the EEG the decoder has to work with is not optimal. Because the EEG is decimated without any anti-aliasing filters (as this would smear out the CI artifacts), it is likely that some additional noise due to aliasing is introduced in the EEG.

The above issues can be alleviated by using EEG acquisition system with high bandwidth, such that the artifact decay time is shortened. This way, gaps can be made shorter, which makes the stimulus less disturbing to listen to. Ultimately, using very high sample rates, sampling could potentially be done in between stimulus pulses, removing the need for gap insertion altogether (unless the artifact decay is partly caused by tissue effects such as volume conduction). This approach would be similar to EASSR measurements, where the samples between pulses of a 500 or 900 Hz pulse train are sampled and used for responses detection (Hofmann & Wouters 2010, Deprez et al. 2017*a*). Another approach might be to exploit the natural gaps in speech rhythms, and use artifact-free EEG during the unstimulated pauses between words.

Another limitation of the study is the use of a linear decoder and relatively simple reference envelopes. As shown in Fig. 8, the introduction of some non-linear processing improves decoding accuracy. However, the simple power law model can be improved to better model the complex auditory processing in the brain and periphery. Another improvement can be made by using more advanced decoding techniques. For instance, the inclusion of more speech features in the decoder, such as spectral or linguistic features, significantly improves decoding results (Di Liberto et al. 2015, Brodbeck et al. 2018). Also non-linear decoding techniques such as neural networks are likely to improve results.

Despite some limitations, this study delivers a strong proof of concept for speech envelope decoding in CI users. The technique can be further improved and refined by using more suitable recording equipment, more sophisticated algorithms and individualized parameters. By enabling the recording and decoding of continuous neural responses to speech in CI users, the auditory and neuroscientific research performed on normal-hearing subjects can be translated to CI users. This will allow to better understand, diagnose and monitor hearing impairment, and will lead to applications that can improve the rehabilitation and listening performance of CI users.

## V. CONCLUSIONS

In the present study, we demonstrated for the first time a technique that allows to assess the neural tracking of the speech envelope in cochlear implant users despite the presence of continuous stimulation artifacts. The technique relies on the insertion of small gaps in the stimulus, which allows sampling of EEG free of artifacts. The technique was validated to be an effective method for eliminated the stimulus artifacts and did not affect the listener’s speech intelligibility.

It was verified that the obtained responses to the speech envelope are of neural origin and that the artifacts do not affect the decoding of the speech envelope. The response decoding can be improved further using subject-specific processing.

The stimulus artifact removal technique may help translate current research with normal hearing persons to severely hearing impaired and deaf populations. Additionally, the demonstrated measure of neural encoding of speech will support future applications based on objective measures for hearing with a cochlear implant.

## Footnotes

1 The stories used were translated fairy tales: “De Wilde Zwanen” (H.C. Andersen) and “Marfoesjka en de vorst”, (Russian fairy tale).

## VI. ACKNOWLEDGEMENTS

This project has received funding from SB PhD grants of the Research Foundation Flanders (FWO) awarded to Ben Somers (1S46117N) and Eline Verschueren (1S86118N), from the KU Leuven Special Research Fund (OT/14/119), and from the European Research Council (ERC) under the European Unions Horizon 2020 research and innovation programme starting grant to Tom Francart (637424).

